# Three-dimensional ATUM-SEM reconstruction and analysis of hepatic endoplasmic reticulum-organelle interactions

**DOI:** 10.1101/2020.11.21.392662

**Authors:** Yi Jiang, Linlin Li, Xi Chen, Jiazheng Liu, Jingbin Yuan, Qiwei Xie, Hua Han

**Affiliations:** National Laboratory of Pattern Recognition, Institute of Automation, Chinese Academy of Sciences, Beijing 100190, China; School of Artificial Intelligence, University of Chinese Academy of Sciences, Beijing 100049, China; School of Future Technology, University of Chinese Academy of Sciences, Beijing 101408, China; Data Mining Lab, Beijing University of Technology, Beijing 100124, China; CAS Center for Excellence in Brain Science and Intelligence Technology, Shanghai 200031, China

**Keywords:** ATUM-SEM, deep learning, liver, ER, MCSs

## Abstract

The endoplasmic reticulum (ER) is a contiguous and complicated membrane network in eukaryotic cells, and membrane contact sites (MCSs) between the ER and other organelles perform vital cellular functions, including lipid homeostasis, metabolite exchange, calcium level regulation, and organelle division. Here, we establish a whole pipeline to reconstruct all ER, mitochondria, lipid droplets, lysosomes, peroxisomes, and nuclei by automated tape-collecting ultramicrotome scanning electron microscopy (ATUM-SEM) and deep-learning techniques, which generates an unprecedented 3D model for mapping liver samples. Furthermore, the morphology of various organelles is systematically analyzed. We found that the ER presents with predominantly flat cisternae and is knitted tightly all throughout the intracellular space and around other organelles. In addition, the ER has a smaller volume-to-membrane surface area ratio than other organelles, which suggests that the ER could be more suited for functions that require a large membrane surface area. Moreover, the MCSs between the ER and other organelles are explored. Our data indicate that ER-mitochondrial contacts are particularly abundant, especially for branched mitochondria. In addition, ER contacts with lipid droplets, lysosomes, and peroxisomes are also plentiful. In summary, we design an efficient method for obtaining a 3D reconstruction of biological structures at a nanometer resolution. Our study also provides the first 3D reconstruction of various organelles in liver samples together with important information fundamental for biochemical and functional studies in the liver.

## Introduction

In all eukaryotes, the endoplasmic reticulum (ER) is a contiguous and complicated membrane network, which is formed by interconnected cisternae and tubules with a single lumen (Zhang and Hu, 2016). The ER extends throughout the cell with a high surface area. In the 1950s, the ER was first identified by observing mouse fibroblasts by electron microscopy (EM, Porter et al., 1945). Historically, the ER is composed of rough ER (ribosome rich) and smooth ER (ribosome free), which are generally organized in cisternae or tubular networks respectively. The polymorphic structure of the ER is intimately related to its many functions, including lipid homeostasis, drug metabolism, secretory protein biogenesis, and regulation of *Ca*^*2+*^ dynamics (Baumann et al., 2001).

The contacts between the ER and other organelles were recognized many years ago (Porter et al., 1957; Rosenbluth, 1962; Csordás et al., 2006). What remained unclear was whether the contacts represented short-term interplays or long-term tethering. For example, membrane contact sites (MCSs) were defined as membrane appositions where the distance between two membrane bilayers was ≤ 30 nm (Wu et al., 2017). The contact is distinguished from vesicle transport and membrane fusion, which are vital for lipid exchange between organelles (Stefan et al., 2017). Subsequent studies focused on the form factors of the MCSs and how they regulate communication between the ER and other organelles (De Brito et al., 2008; Lebiedzinska et al., 2009; Phillips and Votlez, 2016;).

With the rapid development of EM technology, its application in biology has also become widespread (Denk et al., 2004; Knott et al., 2008; Briggman et al., 2012). EM displays the ultrastructure of a region of interest with high resolution (nanometer scale). Furthermore, through 3-dimensional (3D) reconstruction, we can observe their real 3D structure in the cell. Due to the limitation of the low resolution of fluorescence microscopy, the 3D ultrastructure reconstruction of various organelles through EM has become particularly important in the field of biology.

In recent years, the application of the deep-learning technology to biological study has been increasingly investigated (Xiao et al., 2018; Liu et al., 2020). For example, image processing technology has been applied to the contour segmentation of fluorescent protein and EM images. Many previous studies mainly used manual or semimanual methods to obtain the desired 3D reconstruction of the ER and other cell structures with small-scale EM data (Adduda et al., 2014; Wu et al., 2017). To our knowledge, few studies have utilized deep-learning technology for the 3D reconstruction of the ER and other organelles using large-scale EM data.

The liver is one of the most important models for studying the functions of ER-organelle interactions. Here, we initially used automated tape-collecting ultramicrotome scanning electron microscopy (ATUM-SEM) (Briggman et al., 2012) to image liver samples. Then, to map the 3D structure of organelles and their connections within the liver tissue, we applied deep-learning technology to effectively reconstruct all ER, mitochondria, lipid droplets, lysosomes, peroxisomes, and nuclei. Finally, a systematic analysis of the morphology and interactions between the ER and other organelles was presented, which can provide important basic information for biochemical and functional studies.

## Results

We acquired a 3D EM dataset from the liver of an adult C57/BL male mouse by using ATUM-SEM (dataset size: 81.9 × 81.9 × 31.5 *μm*^3^, voxel size: 5 × 5 × 45 *nm*^3^). For the 3D reconstruction (Fig. 2), we first adopted a coarse-to-fine strategy to 3D-align the serial images, and then we obtained a 3D image stack of interest (size: 20 × 20 × 31.5 *μm*^3^, Fig. 2A and Video S1). Afterwards, we designed an image segmentation method based on deep learning to automatically segment the various organelles followed by manual proofreading (See Materials and methods, Fig. S1 and S2). Then, we employed a 3D connection method to calculate the relationships of each organelle in 3D (except for the ER, each organelle is represented by a unique label). We reconstructed all ER, mitochondria, lipid droplets, lysosomes, peroxisomes, and nuclei (the Golgi complex was not reconstructed) to generate a 3D model that mapped the liver samples (Fig. 2A, the number of organelles reconstructed is 3500 for mitochondria, 224 for lipid droplets, 90 for lysosomes, 4035 for peroxisomes, and 7 for nuclei).

**Figure 1.**
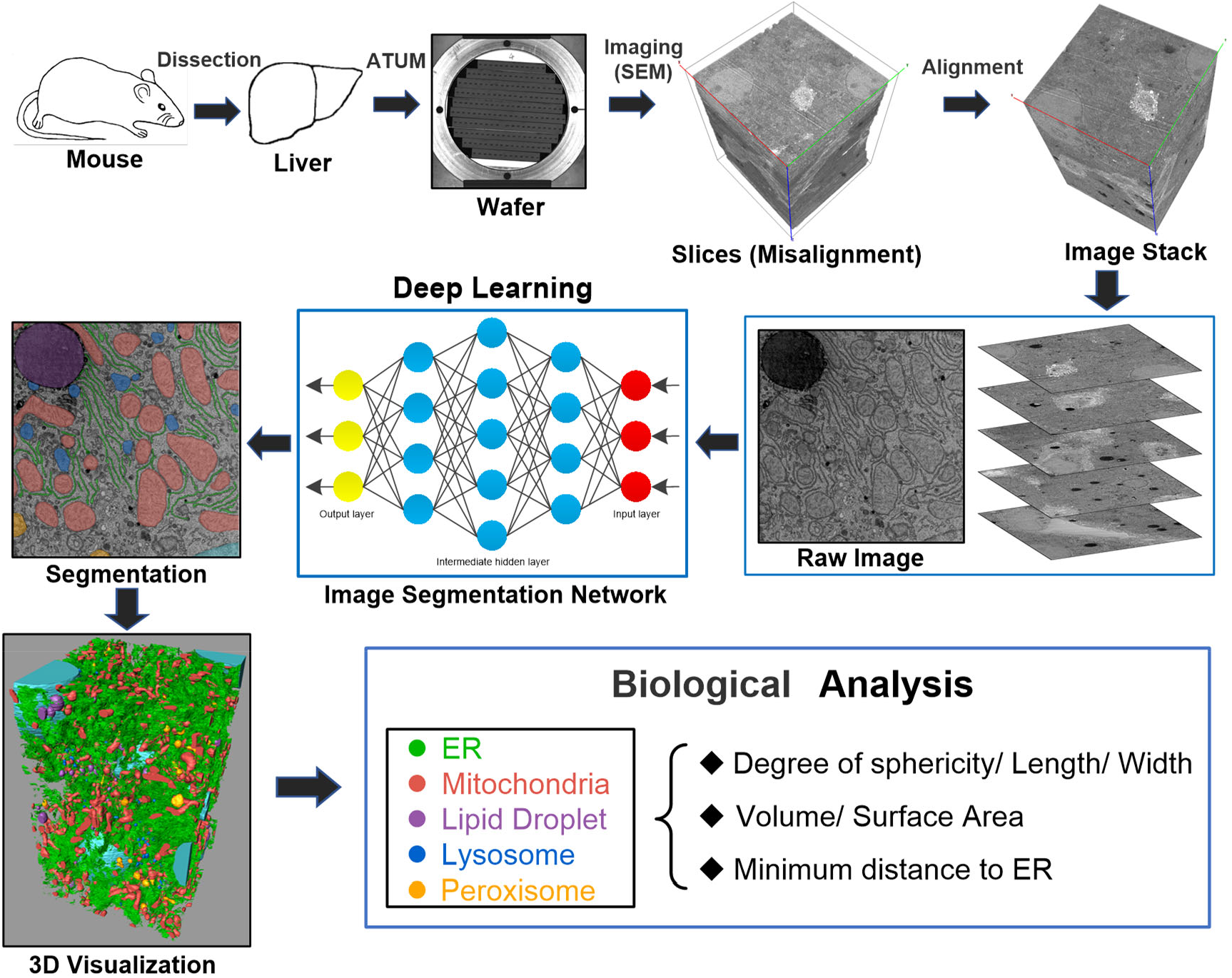
The pipeline of 3D reconstruction and analysis for the mouse liver is based on ATUM-SEM and deep-learning technology. (The first row, left to right) First, the liver of an adult C57/BL male mouse was dissected, then fixed and embedded. After, serial sections of liver samples were continuously cut with the ATUM and collected on tape, the tape was segmented and attached to *Si* wafers. Next, serial sections were imaged by SEM to generate serial images (misalignment). Following, we adopted a coarse-to-fine alignment method to obtain a 3D image stack. (The second row, right to left) All raw 2D EM images were input into the image segmentation network to obtain the segmentation images for the various organelles. (The third row, left to right) 3D visualization provided by Amira software. Afterwards, biological analysis of various organelles based on 3D reconstructions. Note, image segmentation network is a simplified diagram, and please see the supplemental materials for detail (Fig. S1).

**Figure 2.**
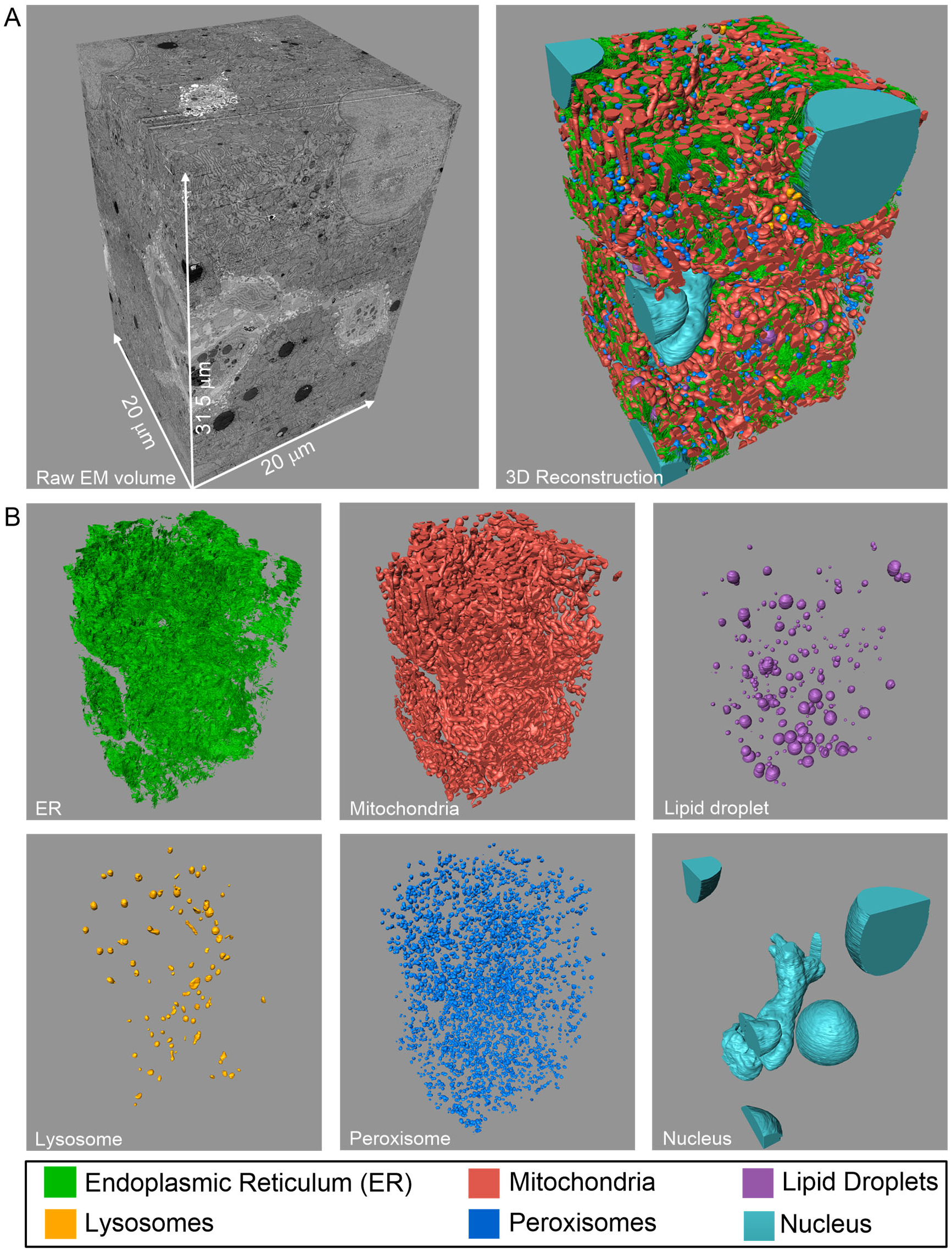
ATUM-SEM volume of mouse liver and 3D reconstruction of various organelles. (A) Raw ATUM-SEM volume of the region of interest from the mouse liver (left), with a corresponding the size of approximately 20 μm × 20 μm × 31.5 μm. The 3D reconstruction of various organelles corresponds to the left image (right). (B) 3D reconstructions of all ER, mitochondria, lipid droplet, lysosome, peroxisome, and nuclei, which correspond to A.

We also visualized the 3D structure of the organelles by using Amira software (Fig. 2B). Furthermore, the morphology and MCSs were systematically analyzed from the 3D model. In Video S2 (Fig. 2B), the fine 3D structure of the ER and other organelles can be observed clearly at the nanoscale, thus minimizing the reliance on techniques that image at low resolution, such as fluorescence imaging, for our potential understanding of these organelles. In addition, the position relationship between the ER and other organelles can also be observed, which provides accurate evidence of whether MCSs are formed between the ER and other organelles.

### A detailed 3D EM structure of the ER and organelles

The 3D EM reconstruction of the liver cell revealed some new information about the structure of the ER. As shown in Video S2 (Fig. 2B), the ER (green) is a contiguous and complicated network that is knitted tightly all over the intracellular space and around other organelles. Additionally, the local morphology of the ER is also diverse. For example, according to the 3D structure, the ER next to the nucleus consists of flat cisternae (Fig. 3A), and some fenestra are observed in the flat ER from the 0° or 180°viewing direction. Most of the fenestrated, flat ER presents with a dense and side-by-side distribution. In addition, local segments of the ER are not independent, but branch out and are interlaced with each other (Fig. 3B). We found that the flat cisternae-shaped exhibits high membrane curvature only at the edge and is wider than the middle section of the lumen (Fig. 3C, white arrow).

**Figure 3.**
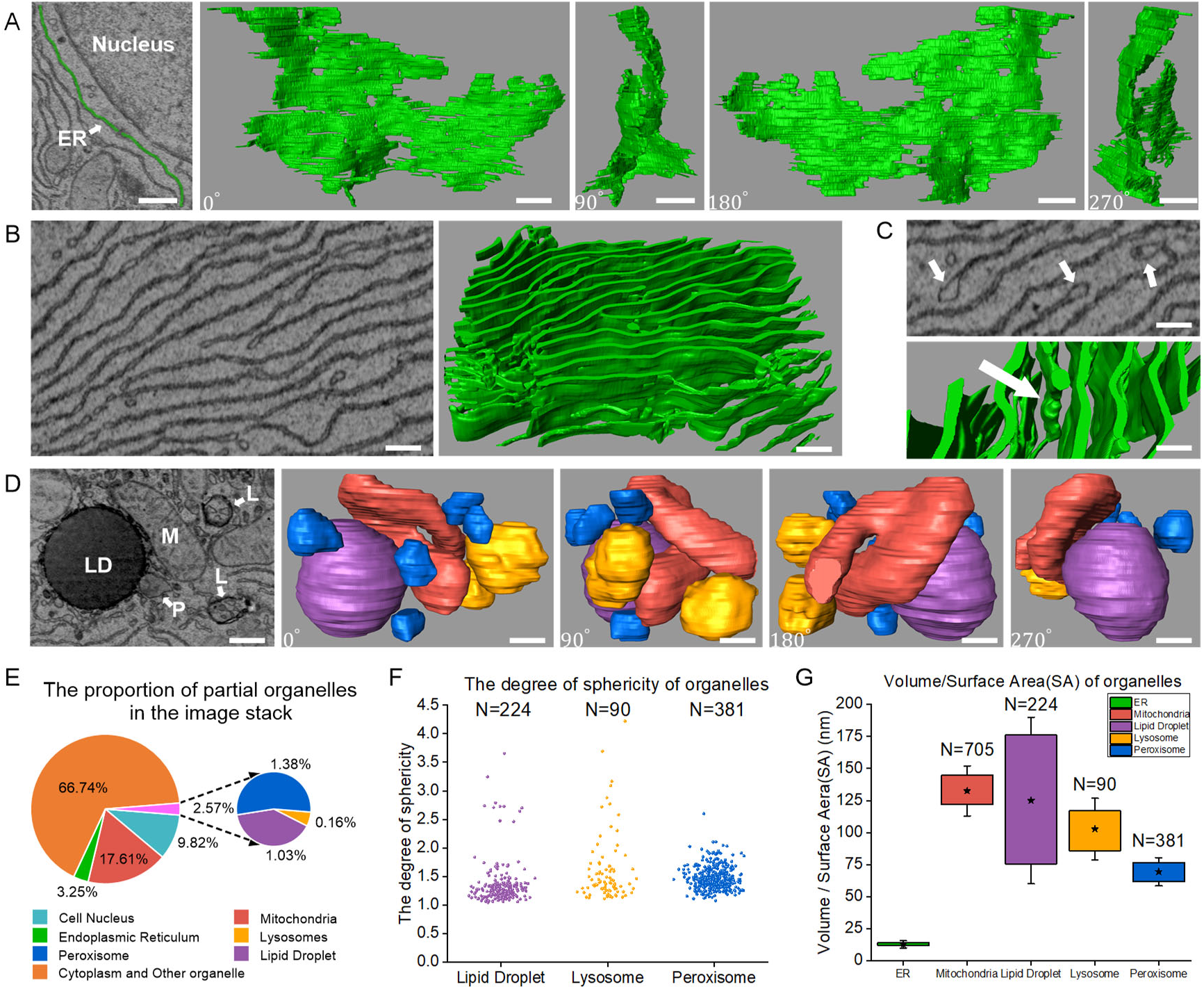
3D ultrastructural analysis of the ER and other organelles morphology. (A) 3D structure of the ER next to the nucleus. The left image is a raw EM image, and the second to fifth images show the 3D structure from different views, followed by the 0°, 90°, 180°, 270° (Clockwise rotation). (B) 3D structure of dense ER area excluding other organelles, which shows branched and interlaced with each other. The left is a raw EM image, the right is a 3D reconstruction. (C) ER presents with high membrane curvature and a wider lumen at the edge (white arrow). Above: raw EM image, below: 3D reconstruction. (D) 3D structure of mitochondria, lipid droplets, lysosomes, and peroxisomes (color-coded as Fig. 2). The first image is a raw EM image, the second to fifth images are 3D reconstructions that were viewed from different perspectives, followed by the 0°, 90°, 180°, 270° (Clockwise rotation). M, mitochondria; LD, lipid droplet; L, lysosome; P, peroxisome. (E) Pie charts show the percent of reconstructed organelles in the liver tissue, as assessed by inspection of ATUM-SEM image stack. (F) Scatter diagram showing the degree of sphericity of lipid droplets, lysosomes, and peroxisomes, the closer to 1, the closer to the perfect sphere (*N*_*lipid droplets*_ = 224, *N*_*lysosomes*_ = 90, *N*_*peroxisomes*_ = 381 measurements were selected randomly). (G) The ratios of volume to the surface area were calculated from 3D reconstruction for ER, mitochondria, lipid droplets, lysosomes, and peroxisomes. ‘*’ shows the means of ratios, and caps show standard error of the mean. (*N*_*mitochondria*_ = 705 measurements were selected randomly, other as F). Scale bars: 750 nm in A; 400 nm in B; 350 nm in C; 450 nm in D.

As depicted in Fig. 3D, the volume of a single lipid droplet is large, and the volume of a single peroxisome is the smallest. In addition, mitochondria present with an irregular shape on direct visual inspection. The nucleus was also reconstructed in the image stack. Fig. 3E shows the volume ratios of the reconstructed organelles in the whole tissue. Obviously, the cytoplasm is the largest part of the volume. Among the reconstructed organelles, the total volume of mitochondria is the largest. These results indirectly indicate that the energy supply from mitochondria is essential for liver cells. Surprisingly, although the ER is dense and consists of numerous segments (Video S2), the volume occupied by the ER is only 3.25%. Because the double membranes of the ER are close together, a narrow lumen is formed. In addition, lipid droplets and lysosomes are relatively uncommon.

As seen from Fig. 3D, the lipid droplets, lysosomes, and peroxisomes are approximately spherical, so their sphericity was measured for the entire volume (Fig. 3F). The closer the sphericity is to 1, the more it tends to be a perfect sphere. A total of 224 and 90 lipid droplets and lysosomes, respectively, were found; we randomly selected 381 to be measured among all reconstructed peroxisomes. The scatter plot shows that most of the lipid droplets and peroxisomes tend to be spherical, while lysosomes tend to beellipsoidal.

We used our 3D reconstruction to calculate the volume-to-surface area ratios (V/SA) of all reconstructed organelles (Fig. 3G). A total of 705 of the 3500 mitochondria were randomly selected for statistical analysis; these mitochondria were intact and did not appear at the boundary of the volume. The data revealed that the volume-to-membrane surface area ratios of various organelles are distinguishable. In the liver, the ER presents with a smaller volume-to-membrane surface area ratio than the other organelles, which suggests that the ER could be better suited for functions that require a large membrane surface area, whereas mitochondria, lipid droplets, lysosomes, and peroxisomes may be more suitable for functions that need more internal space. For instance, mitochondria require a large amount of internal space to form the complex mitochondrial cristae structure.

### Mitochondria are unique organelles in the liver

Fig. 3E shows that mitochondria are one of the most abundant organelles in the liver. Mitochondria play an important role in the homeostasis of carbohydrate, lipid, and protein metabolism in many organisms (McBride et al., 2006). The morphology of the mitochondria in the liver is diverse. Fig. 4A and Fig. 4B show unbranched and branched mitochondria. In addition, it can be observed that the size of the mitochondria is also different from that shown in Video S2. Branched mitochondria may be related to division and fusion, because mitochondrial proliferation and division are common phenomena in liver cells. However, we also cannot rule out the possibility that branched mitochondria are original shapes and are not related to division and fusion.

**Figure 4.**
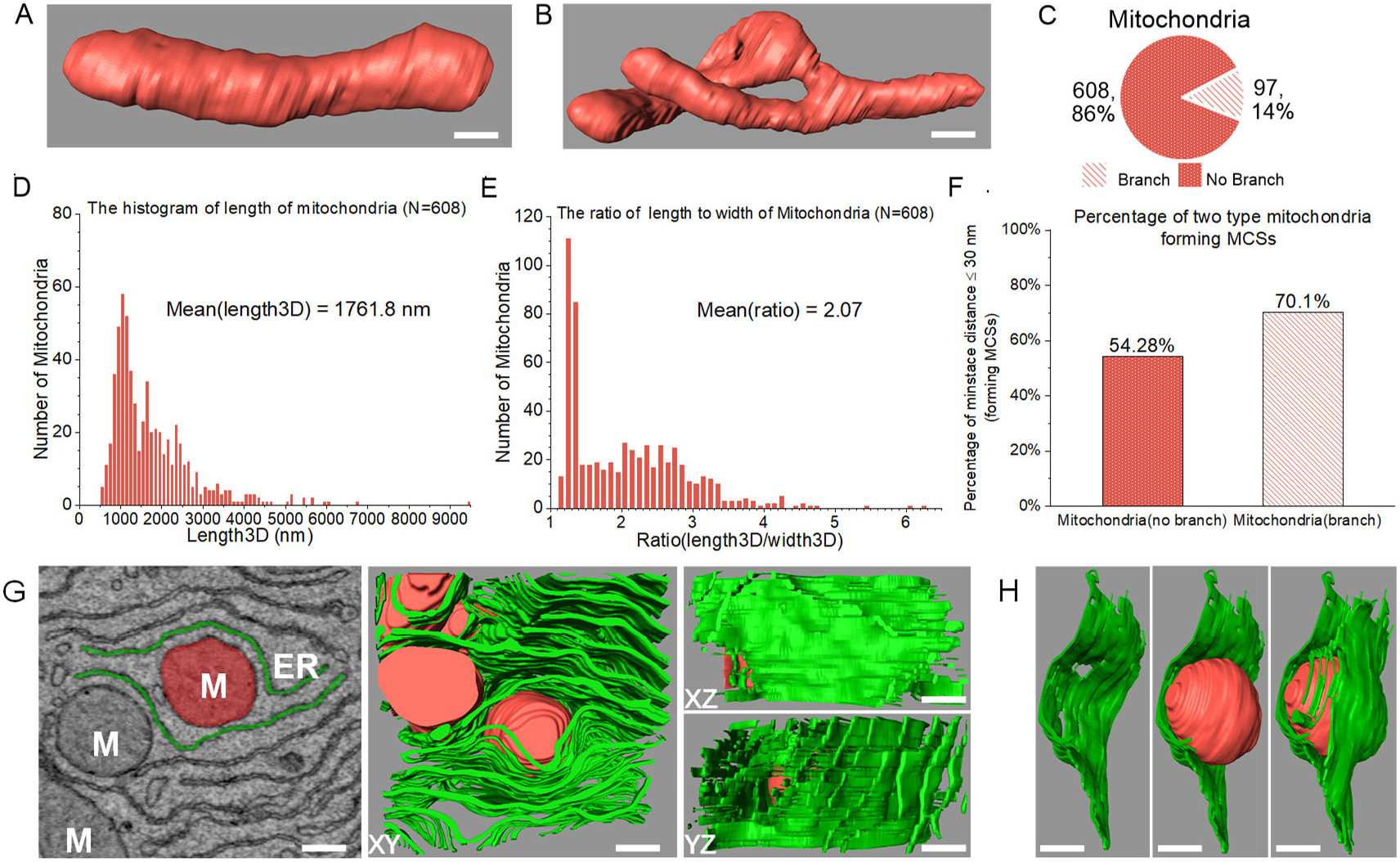
3D structural analysis of mitochondria in the liver. (A) Unbranched mitochondria. (B) Branched mitochondria. (C) Pie chart shows the percent of unbranched and branched mitochondria (total 705, as Fig. 3G. The shape of all mitochondria is intact). (D) The histogram shows the length of unbranched mitochondria in 3D reconstruction, length3D mean = 1761.8 nm (N = 608, as C). (E) The histogram shows the ratio of length to width of unbranched mitochondria in 3D reconstruction, ratio mean = 2.07 (N = 608, as C). (F) The bar chart shows the percent of two types of mitochondria forming MCSs. Percentages are indicated above each graph. (G) The spatial relationship between mitochondria and ER. The first figure shows a raw EM image, the other figures show the 3D structure from XY, XZ, and YZ perspectives respectively (M, mitochondria). (H) Mitochondria are surrounded by the ER, as G. Scale bars: 250 nm in A; 350 nm in B; 190 nm in G, 280 nm in H.

In Fig. 4C, we found that unbranched mitochondria are predominant. The length of the unbranched mitochondria was then measured in the 3D reconstruction, revealing that most ranged from 1000 nm ~ 2500 nm (mean = 1761.8 nm). Surprisingly, the longest mitochondria exceeded 9000 nm (Fig. 4D). The ratio of length to width of the mitochondria can be used as a reference to judge abnormalities. Our results show that this ratio is generally less than 3 (mean = 2.07), and very few ratios exceed 6 (Fig. 4E). After visualization of the spatial relationship between the mitochondria and the surrounding ER (Fig. 4G, H), we observed that mitochondria are wrapped in the ER, which is a common phenomenon in the liver. This observation was not made by chance, which means that there are crucial connections between the mitochondria and the surrounding ER.

### MCSs are formed between the ER and the mitochondria, lipid droplets, lysosomes, and peroxisomes

Many previous studies have shown that MCSs are essential for the exchange of biological substances between the ER and other organelles. These two membranes contact but do not fuse, delivering substances in the form of non-vesicles. Recently, ER-mitochondria contacts have become a hotspot of research. The ER engages with mitochondria at specialized ER domains known as mitochondria-associated membranes (MAMs), which are indispensable for mitochondrial dynamics and function (Zhou et al., 2020).

Obviously, the MCSs between mitochondria and ER were observed easily when we examined the raw 2D EM images (Fig. 5A, left). The white arrow indicates the formation of MCSs between the mitochondria and ER. The second to fourth images (Fig. 5A) show the 3D reconstruction from different views (XY, XZ, and YZ directions). Meanwhile, we measured the minimum distance between the mitochondria and the surrounding ER in the 3D reconstruction. According to the sampling survey statistics, we randomly extracted 949 from all 3500 reconstructed mitochondria. Fig. 5E (red dot on the left) shows the distribution of the minimum distance between the 949 mitochondria and the surrounding ER. Through qualitative observation, the minimum distance between most mitochondria and the ER is within 60 nm. The mean minimum distance was calculated (Fig. 5F, the red bar on the left). To determine how many mitochondria and ER form MCSs (minimum distance ≤ 30 nm), we computed the percentage of mitochondria that form MCSs (Fig. 5G, the red bar on the left). Approximately 53.7% of mitochondria and ER formed MCSs.

**Figure 5.**
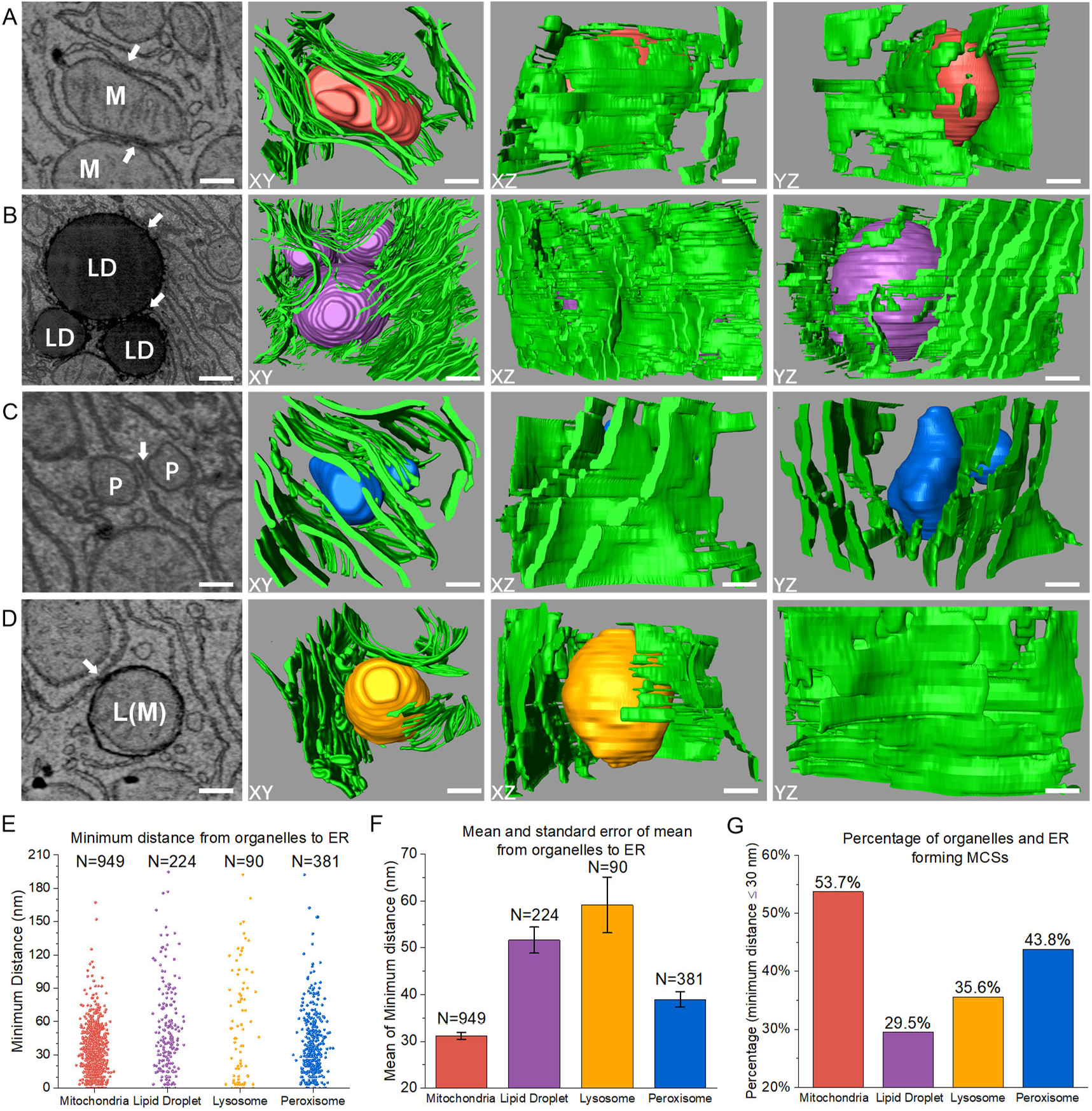
A 3D analysis of membrane contact sites (MCSs) between the ER and other organelles. (A) MCSs are formed between the ER and mitochondria. The first image shows the raw EM image, the white arrow indicates MCSs in a 2D image. The second to fourth images show that the 3D reconstruction is based on the first image, which was viewed from different perspectives (followed by XY, XZ, and YZ directions). (B-D) MCSs are formed between ER and lipid droplets, lysosomes, and peroxisomes, respectively (illustration as A). (E) Scatter diagram showing the minimum distance between organelles and ER. MCSs are formed within 30 nm. *N*_*mitochondria*_ = 949 measurements, other as Fig. 3F. (F) Bar chart showing the mean of the minimum distance. The caps show a standard error of the mean. (G) Bar chart showing the percent of organelles forming MCSs with ER. Percentages are indicated above each graph. Color-coded as Fig. 2. Scale bars: 290 nm in A; 550 nm in B; 210 nm in C; 260 nm in D.

When examined in cross-section, the ATUM-SEM images show the position of lipid droplets and the surrounding ER. In Fig. 5B (left), the white arrows indicate the MCSs between lipid droplets and the ER. The second to fourth images (Fig. 5B) show the 3D reconstruction viewed from different perspectives (XY, XZ, and YZ directions). The distribution of the minimum distance between lipid droplets and the surrounding ER is shown in Fig. 5E (purple dot in the middle). The mean minimum distance is 50 nm (Fig. 5F, purple bar in the middle). The percentage of lipid droplets that formed MCSs with the surrounding ER was approximately 29.5% (Fig. 5G).

Similarly, we observed the structure of lysosomes and peroxisomes in the 2D EM image, with the white arrows indicating the MCSs (Fig. 5C, D). Likewise, their 3D reconstruction was viewed from different perspectives (XY, XZ, and YZ directions). We found that the minimum distance between lysosomes and the ER was not clustered but was similar to a random distribution. However, the minimum distance between most peroxisomes and the ER is less than 60 nm (Fig. 5E). The mean minimum distances between lysosomes and peroxisomes and the ER were 60 nm and 40 nm, respectively (Fig. 5F). Approximately 35.6% of lysosomes and 43.8% of peroxisomes form MCSs with the ER (Fig. 5G).

## Discussion

For the first time, we established a whole pipeline to reconstruct the ER and other organelles in the liver by ATUM-SEM (Fig. 1), which generated an unprecedented 3D model to map the liver sample. Furthermore, the 3D model allowed a systematic analysis of the distribution and abundance of various organelles and a quantification of the MCSs between the ER and other organelles. In our workflow, ATUM-SEM was adopted to generate image data because it is suitable for large volumes of data, and serial sections can be imaged again when the first imaging fails. In addition, the pixel resolution of ATUM-SEM (x- and y- directions, 5 nm × 5 nm) was sufficient for recognizing the membranes of the ER and other organelles. The main drawback of the ATUM-SEM technique is the low resolution in the z-direction (45 nm), which leads to the inability to visualize some tissues. However, for biological structures over a hundred nanometers in size, the impact can be ignored. For 3D reconstruction, the ATUM-SEM method requires a more accurate alignment technology to align the serial images than FIB-SEM in situ imaging.

To our knowledge, we are the first to apply deep-learning technology to automatically segment the ER and other organelles in a 3D EM image stack. Compared with previous semimanual methods, we have a greater advantage in processing large-scale data. For example, the membrane contours of the ER and other organelles were traced semimanually by using 3Dmod software (West et al., 2011, Wu et al., 2017). In addition, our method of segmentation for various organelles is also applicable to FIB-SEM data, provided that the data are reliable. Therefore, our reconstruction workflow can efficiently obtain the 3D structure of various organelles so that biologists can focus more on the study of biological function.

Our 3D model revealed that the ER is distributed densely throughout the tissue to form a continuous membrane network and that the local morphology is diverse. We observed that the ER commonly presents with flat cisternae, and the lumen between membrane bilayers is narrow. It was somewhat surprising to us that the fenestrations and edges have a higher membrane curvature than the middle sections in the flat cisternae and that the lumen of the cisterna edges is wider (Fig. 3C). As reported previously, reticulons (Rtns) and Yop1 proteins are related to the membrane curvature of ER, and the deletion of these two proteins leads to a loss of cisternal fenestrations (West et al., 2011). We did not perform relevant experiments to confirm the abundance of the two proteins in these regions. Our data are not suitable for reconstructing the tubular ER because it is difficult to distinguish between tubular ER and free vesicles in the 2D EM images. However, we can observe that the cisternal ER is very dominant in the original EM image stack (Video S1).

The unprecedented 3D model shows the spatial distribution and abundance of various organelles (Video S2; Fig. 2A, B). In addition to the ER, mitochondria have the tightest distribution and diverse forms. Mitochondria continuously divide and fuse to form a tight network in liver cells. However, the mitochondria are static in our 3D model because they were imaged by EM at a specific moment. Therefore, we cannot ensure whether branched mitochondria are dividing, fusing, or maintain their original shape. Previous studies have suggested that the abnormal sizes of some mitochondria were related to some liver diseases. For example, giant mitochondria in hepatocytes were related to alcoholic liver disease (Bruguera et al., 1977).

In recent years, the connection between the ER and other organelles has been studied extensively in eukaryotic cells. Increasing numbers of researchers have realized the importance of the MCSs in cell physiology. Universally, MCSs are studied by fluorescence images or 2D ultrastructures. Here, we used ATUM-SEM to generate a 3D model to analyze the relationship between the ER and other organelles. Due to the static imaging of EM, we should be cautious about the findings that we observe from the 3D structure. However, most cells in the liver should have common features, so our results can provide basic information for biochemical and functional studies of MCSs.

Although the ER and mitochondria play different roles in cells, their interaction is necessary for the exchange of calcium, lipids, and metabolites (Flis et al., 2013). Many previous studies have focused on the MCSs between the ER and mitochondria, which are defined as MAMs. Here, we found that more than half of the mitochondria formed MAMs with the ER from the 3D reconstruction. Moreover, branched mitochondria are more likely to form MAMs with the ER than unbranched mitochondria. Remarkably, branched mitochondria may be in the process of division, and previous studies have indicated that ER-mitochondrial contacts coordinate mtDNA replication with mitochondrial division in yeast and human cells (Murley et al., 2013; Lewis et al., 2016). Additionally, a previous study showed that obesity can lead to a marked reorganization of MAMs resulting in mitochondrial calcium overload in the liver, compromising mitochondrial oxidative capacity and augmenting oxidative stress (Arruda et al., 2014).

Lipid droplets (LDs) are storage organelles for neutral lipids (Wu et al., 2018). Contacts between the ER and lipid droplets are frequent and conspicuous. However, our data show that the lipid droplet is not enfolded by the ER; instead the edge of the ER is in contact with and displays continuity with the LD membrane, displaying the continuity of the membrane (Fig. 5C). These contacts have been defined as membrane bridges, which are closely related to the unique structure and biological process of LDs.

Lysosomes are essential organelles in the cell that can decompose substances that enter the cell from the outside world and digest the local cytoplasm and organelles of the cell itself. Recent study has shown that ER-lysosome contacts are signaling hubs that enable cholesterol sensing by mTORC1, and targeting the sterol-transfer activity of these signaling hubs could be beneficial in patients with Niemann-Pick disease, type C (NPC) (Lim et al., 2019). Our data show that the lysosome is low density and that it can take on a variety of shapes, such as spherical, ellipsoidal, and elongated. In addition, we observed that lysosomes dissolve mitochondria and peroxisomes. Furthermore, the MCSs between the ER and lysosomes are also indispensable to the physiological function of cells.

Peroxisomes are ubiquitous organelles in cells. The peroxisome plays a crucial role in metabolism, and many of these metabolic functions are carried out in partnership with the ER (Wu et al., 2018). In addition to the ER, peroxisomes are intimately associated with mitochondria and lipid droplets, and their ability to carry out fatty acid oxidation and lipid synthesis, especially the production of ether lipids, may be critical for generating cellular signals required for normal physiology (Lodhi et al., 2014). However, peroxisomes have received little attention due to their small size. Our data show that peroxisomes are the smallest organelles in our reconstructed image stack but are omnipresent (Video S2, Fig. 2B). Like lipid droplets, peroxisomes are also considered to be derived from the ER. Contact between peroxisomes and the ER is also frequent (Fig. 5D).

In summary, our results enhance the knowledge about the 3D structure, distribution, and abundance of the various organelles in liver cells. They also enable us to determine the MCSs between the ER and other organelles at a nanometer resolution. However, our results only present qualitative and quantitative information from a structural perspective based on static imaging. Therefore, our findings require further EM and dynamic imaging to verify their uniqueness, including whether the MCSs between the ER and other organelles represent short-term interplay or long-term tethering and whether the same phenomenon occurs in other tissues of the liver. Notably, MCSs are known to be linked to liver diseases. The ablation of Mfn2 in the liver newly revealed that the destruction of ER-mitochondrial phosphatidylserine (PS) transfer is one mechanism involved in the development of liver disease (Hernández et al., 2019).

## Materials and methods

### Animals

An adult C57/BL male mouse was kept in a temperature-controlled room with a 12-hour light-dark cycle, and we fed it standard mouse chow and normal water. All animal care procedures and research were approved by the Institutional Animal Care and Use Committee of Chinese Academy of Science.

### Fixation of mouse liver tissue

The mouse liver tissues were immersed in 4% (w/v) PFA and 2.5% (Sigma, G5886) GA. Then, the samples were fixed in 2% OsO4 (Ted Pella, 18451) in phosphate buffer (0.1M, pH7.4) for 90 mins at room temperature. Then, the staining buffer was replaced with 2.5% ferrocyanide (Sigma, 234125) in phosphate buffer (0.1 M, pH 7.4) for another 90 mins at room temperature. The samples were washed 3 times with 0.1 M phosphate buffer and incubated with filtered thiocarbohydrazide (TCH, Sigma, 223220) for 45 mins at 40 °C. Next, the samples were fixed again with unbuffered 2% OsO4 for 90 mins, and then incubated overnight in 1% uranyl acetate aqueous solution at 4 °C. After the samples were incubated with a lead aspartate solution (0.033 g lead nitrate (Sigma, 228621) dissolved in 5 ml 0.03 M aspartic acid (Sigma, 11189, pH 5.0)) for 120 mins at 50 °C. We dehydrated the sample through a gradient ethanol series (50, 70, 80, 90, and 100% ethanol, 10 mins each) and pure acetone. Finally, the samples were embedded with Epon 812 resin (SPI, 02660-AB).

### ATUM-SEM imaging

Serial sections of liver samples were continuously cut through a commercial ATUM with a diamond knife (Diatome, MC16425) and collected on Kapton polyimide tape (width 8 mm, thickness 100 μm). Then, the tape was segmented and attached to 4-inch silicon wafers by a double-coated carbon conductive tape (Ted Pella). Next, the wafers were coated with 6 nm of carbon via a high vacuum film deposition instrument (Leica) to avoid charging. Eventually, serial sections were imaged by scanning electron microscopy (SEM, Zeiss Gemini 300) with a resolution of 5 nm/pixel and a dwell time of 2-5 μs.

### Image alignment

After the serial image dataset was acquired through SEM, all images were inspected manually. If sections were missed or contaminated during imaging, the sections could be imaged again. Since we adopted the ATUM method and the samples were not imaged in situ, nonlinear distortion of the images was inevitable. Therefore, serial image alignment was indispensable in obtaining a 3D EM image stack. In this case, we adopted a coarse-to-fine strategy to align the serial images. First, we performed coarse alignment by extracting corresponding points between adjacent sections with an affine transformation model on the raw serial images (16000 × 16000 pixels). Then, we intercepted the region of interest (6000 × 6000 pixels). Next, we performed fine alignment (Chen et al., 2018), which involves pairwise correspondence extraction between adjacent sections by SIFT-flow (Liu et al., 2010), correspondence position adjustment for all sections, and image wrapping by the moving-least-square (MLS, Schaefer et al., 2006) method. Because the image will contain a small offset to produce black borders after fine alignment, we intercepted the area without the black border (4000 × 4000 pixels) through the entire serial image stack. Thus, we acquired a stack of 702 images (4000 × 4000 pixels), with a corresponding size of approximately 20 μm × 20 μm × 31.5 μm.

### Workflow for 3D reconstruction

The workflow for 3D reconstruction of the 3D EM image stack was as follows. We first designed an image segmentation network to automatically segment all ER, mitochondria, lipid droplets, lysosomes, peroxisomes, and nuclei (Fig. S1). Each raw 2D EM image was input to the deep learning network to obtain the segmentation images for the various organelles. The performance of our image segmentation network is presented quantitatively (Dice coefficient: 0.882 for the ER, 0.9797 for mitochondria, 0.9835 for lipid droplets, 0.9213 for lysosomes, 0.9064 for peroxisomes, and 0.9887 for the nucleus, see Tab. S1) and qualitatively (Fig. S2). Next, we employed a 3D connection method (Li et al., 2018) to calculate the relationship between each organelle in 3D (except for the ER, each organelle is represented by a unique label). For the architecture and implementation details of the image segmentation network, see the supplemental materials.

### 3D visualization and quantification

Amira software (Stalling et al., 2005) was used for 3D visualization of the ER and other organelles. In addition, some morphological measurements of the organelles were obtained through Amira software, such as 3D volume, 3D surface area, 3D length, and sphericity (see supplemental materials). Furthermore, the minimum distance between the ER and other organelles was obtained by measuring the closest distance between their contours in the 3D model.

## Compliance and ethics

The authors declare that they have no conflict of interest.

## Acknowledgements

This work was supported by Special Program of Beijing Municipal Science & Technology Commission (No. Z181100003818001, No. Z181100000118002) and the Strategic Priority Research Program of Chinese Academy of Science, Grant No. XDB32030200 and Bureau of International Cooperation, CAS (No. 153D31KYSB20170059) and International Partnership of Chinese Academy of Science, Grant No. 153D31KYSB20170059. We would like to thank Lixin Wei, Hongtu Ma, and their colleagues (Institute of Automation, CAS) for their assistance in electron microscopy. We extend a special thanks to Chi Xiao and Weifu Li for critical reading and editing of the manuscript. We also thank the Core Facilities of Life Science, Peking University for assistance with Amira software work.

## Author contributions

Yi Jiang, Linlin Li, Qiwei Xie, and Hua Han designed the research; Yi Jiang and Linlin Li performed the research; Linlin Li performed ATUM-SEM imaging; Xi Chen performed the serial images alignment; Yi Jiang performed 3D reconstruction; Yi Jiang, Jingbin Yuan, and Jiazheng Liu performed 3D visualization; Yi Jiang and Linlin Li analyzed and interpreted the data; Yi Jiang wrote the manuscript.

## Supplemental materials

**Video S1.** 3D ATUM-SEM image stack.

**Video S2.** 3D reconstruction of the ER, mitochondria, lipid droplets, lysosomes, peroxisomes, and nuclei in the raw 3D EM image stack.

**Figure S1.**The architecture of our designed image segmentation network.

**Figure S2.** The segmentation results of our designed image segmentation network.

**Table S1.** Dice coefficient of the various organelles.

Technical details of workflow for 3D reconstruction are described, including description of image segmentation network, network training, performance evaluation, proofreading, and description of measurements.

### Captions for movies

**Video S1. 3D ATUM-SEM image stack.**

Serial SEM images (702 sections, 45 nm/interval) from the adult C57/BL male mouse liver. The pixel resolution is 5nm. The size of the raw EM image is 4000 × 4000 (pixel). The region of interest is 20 μm × 20 μm (Here, images were downsampled twice to generate a video).

**Video S2. 3D reconstruction of ER, mitochondria, lipid droplets, lysosomes, peroxisomes, and nuclei.**

This video rotates clockwise 8 times. The first time showing a 3D ATUM-SEM image stack, the size of the sample is 20 μm × 20 μm × 31.5 μm. The second time showing the 3D reconstruction of various organelles (ER, mitochondria, lipid droplets, lysosomes, peroxisomes, and nuclei). For the next six times showing reconstruction of lipid droplets, lysosomes, peroxisomes, mitochondria, nuclei, and ER in sequence.

## Methods

### Image segmentation network

We carefully observed the structure of the various organelles on the 2D EM image. Due to the tangential problem in the cutting of the tissue block, the various organelles showed scale diversity in the 2D EM image. Therefore, we designed a deep convolution neural network with encoder and decoder modules to segment the various organelles in the 2D EM image, which effectively addressed the problem of biological structures segmentation. The architecture of our designed network was illustrated in Fig. S1.

**Figure S1.**
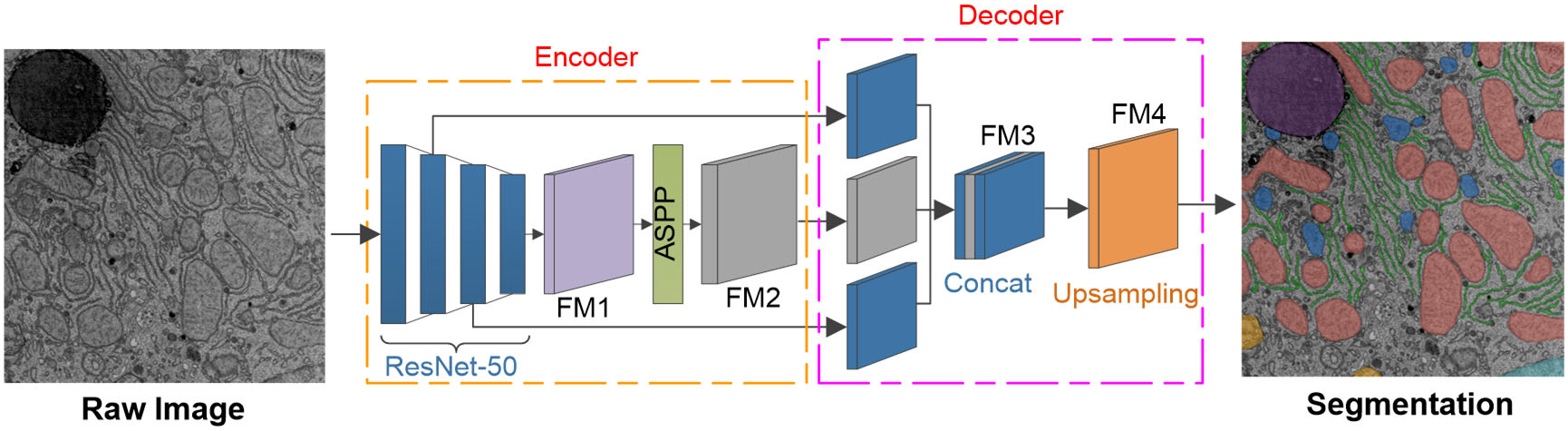
The architecture of our designed image segmentation network. The network contains an encoder module and a decoder module. The raw EM image was fed into the encoder module, which contains ResNet-50 and ASPP. After, three-level features were inputted into decoder module, including Concat and Upsampling. Finally, the output of the network is the segmentation of various organelles.

### Encoder module

We adopted modified ResNet-50 (He et al., 2016) as a network backbone to extract image features. To increase the receptive field of the convolution operation and avoid excessive reduction of the feature map resolution, we used atrous convolution (Chen et al., 2017) in block 4 of the ResNet-50. Note that the output stride is the ratio of the input image spatial resolution to the final output feature map resolution. The output stride is 32 in the original ResNet-50. As we all know, the lower part of the network learns texture information and the upper part learns semantic information, thus a feature map is difficult to get the rich texture and semantic information without any intelligent operation. In addition, the stride is 2 between the adjacent blocks in the original ResNet-50. As confirmed by the empirical experiments, consecutive striding is harmful for semantic segmentation due to the decimation of the detailed information. Here, we applied atrous convolution instead of stride in block 4, thus the output stride is 16 in the modified ResNet-50.

When generating serial sections of biological tissue, the 3D organelles were presented by a multi-scale 2D section. To obtain multi-scale contextual information of organelles, we employed the Atrous Spatial Pyramid Pooling (ASPP, Chen et al., 2018) to process the feature map from modified ResNet-50. Next, we concatenated five feature maps from ASPP and fed the feature map to a 1 × 1 convolution with 256 filters. In closing, the output of the encoder module is a feature map that contains 256 channels of rich semantic information.

### Decoder module

From the above, the feature map resolution from the encoder module is 1/16 of the input image (output stride = 16). Our decoder module combines low-level high-resolution features with high-level low-resolution features, which includes three branches of feature maps. To be specific, the first branch is high-level features from the encoder output, and it is upsampled by a factor of 4. The second branch is low-level features from the output of the block1 in the modified ResNet-50, and then we adopted a 1 × 1 convolution to reduce the number of channels. The final branch comes from the output of the block2, and we applied another 1 × 1 convolution to reduce the number of channels, then the output is upsampled by a factor of 2. After the concatenation, we applied two 3 × 3 convolutions with 256 filters and fed the result to 1 × 1 convolution with 7 filters followed by another simple bilinear upsampling by a factor of 4. The decoder module can recover the detail information of the object and reduce the computational complexity.

Eventually, given an EM image, after the encoder module and the decoder module, a segmentation image of various organelles with the same size as the input image is output.

### Network training

#### Training dataset

To create a dataset for segmentation of the various organelles, we extracted 25 images from 702 images (interval 30 images) because of the high similarity between adjacent sections images for the EM image stack. The ground truth masks of various organelles were manually annotated by using the software tool TrakEm2 (Cardona et al., 2012) for 25 images. We used 20 images as the training data, and the remainder as testing data. Due to GPU memory constraints, the network input size was set to 512 × 512. Here, we divided each image (4000 × 4000, pixel) into 64 images (512 × 512, pixel), the overlap of the width and height are 162 and 108 respectively. In this case, we got 1280 images as training data and 320 images as testing data.

#### Data Augmentation

Overfitting is a common problem in deep learning, mainly caused by two reasons: little data and complicated network. After our network was designed, we adopted a data augmentation strategy to avoid the overfitting of the network. By rotating and folding each image, the number of images was expanded by 8 times, so we got 10240 training images (512 × 512, pixel). The testing data did not perform data augmentation. These images are sufficient for network training. Then, we randomly divided the training data into a training set and a validation set, with 7168 images in the training set and 3072 images in the validation set.

#### Training and testing

The network was trained on the training sets by using the Keras deep learning library and TensorFlow backend (Kreas, 2015). In the training process, we adopted a two-stage training strategy. First training 100 epochs with a learning rate of 0.0001, and then training 50 epochs with a learning rate of 0.00001. Besides, our network was optimized by Adaptive Moment Estimation (Adam) with the following optimization hyperparameters: *β*_1_ = 0.9, *β*_2_ = 0.999 and *epsilon* = 1*e* − 8 for numerical stability. Meanwhile, we used *Cross Entropy* as loss function. It took nearly 24 hours to train our network with the batch size of 4 on a server equipped with an Intel i7 CPU, 512 GB of main memory, and a V100 GPU. After, we tested the performance of our network on the test set.

### Performance evaluation

#### Evaluation criteria

To quantitatively evaluate the quality of pixel-level segmentation, we selected *Dice coefficient* as evaluation criteria, which is a common criterion in biological image segmentation tasks.

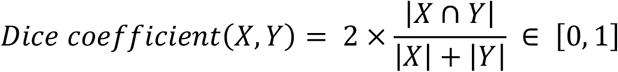

It computes the similarity between the ground truth (Y) and the segmentation result (X). Here, a larger dice coefficient means better segmentation results.

#### Quantitative

To demonstrate the effectiveness of our designed images segmentation network, we presented the quantitative assessment of various organelles (Tab. S1).

**Table S1.** Dice coefficient of the various organelles.

| Organelles | ER | Mitochondria | Lipid Droplet | Lysosome | Peroxisome | Nucleus |
| --- | --- | --- | --- | --- | --- | --- |
| Dice coefficient | 0.8820 | 0.9797 | 0.9835 | 0.9213 | 0.9064 | 0.9887 |

#### Qualitative

To intuitively evaluate our designed image segmentation network, we presented the qualitative assessment of the various organelles (Fig. S2).

The above quantitative and qualitative evaluation shows the image segmentation network is capable of processing large-scale data. Besides, it got rid of the inefficient semi-manual method, which draws the contour of the organelles manually by software.

#### Proofreading

Due to the ATUM-SEM method, image pollution is difficult to avoid completely. The network cannot obtain the precise segmentation of organelles in the contaminated area. Therefore, in order to get an accurate 3D reconstruction, we proofread the contaminated area based on segmentation.

### Description of measurements

Detailed description of the measurements are as follows.

3*D surface area*: Area of the object boundary.

3*D volume*: Volume of the object.

3*D length*: Maximum of Feret Diameters of the object.

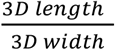: 3*D length* is the maximum of Feret Diameter and 3*D width* is the minimum of Feret Diameter in the orthogonal.

*Sphericity*: 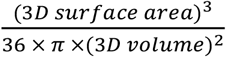, which equals 1 for a perfect sphere.

## References

Zhang, H. and Hu, J. (2016). Shaping the endoplasmic reticulum into a social network. Trends in cell biology, 26(12), 934–943.

Porter, K.R., Claude, A. and Fullam, E.F. (1945). A study of tissue culture cells by electron microscopy: methods and preliminary observations. The Journal of experimental medicine, 81(3), 233.

Baumann, O. and Walz, B. (2001). Endoplasmic reticulum of animal cells and its organization into structural and functional domains. International review of cytology, 205, 149–214.

Porter, K. R., and Palade, G. E. (1957). Studies on the endoplasmic reticulum: III. Its form and distribution in striated muscle cells. The Journal of Cell Biology, 3(2), 269–300.

Rosenbluth, J. (1962). Subsurface cisterns and their relationship to the neuronal plasma membrane. The Journal of cell biology, 13(3), 405–421.

Csordás, G., Renken, C., Várnai, P., Walter, L., Weaver, D., Buttle, K. F., Balla, T., Mannella, C. A., and Hajnóczky, G. (2006). Structural and functional features and significance of the physical linkage between ER and mitochondria. The Journal of cell biology, 174(7), 915–921.

Wu, Y., Whiteus, C., Xu, C. S., Hayworth, K. J., Weinberg, R. J., Hess, H. F., and De Camilli, P. (2017). Contacts between the endoplasmic reticulum and other membranes in neurons. Proceedings of the National Academy of Sciences, 114(24), E4859–E4867.

Stefan, C. J., Trimble, W. S., Grinstein, S., Drin, G., Reinisch, K., De Camilli, P., Cohen, S., Valm, A. M., Levine, T. P., Iaea, D. B., et al. (2017). Membrane dynamics and organelle biogenesis-lipid pipelines and vesicular carriers. BMC biology, 15(1), 1–24.

De Brito, O.M. and Scorrano, L. (2008). Mitofusin 2 tethers endoplasmic reticulum to mitochondria. Nature, 456(7222), 605–610.

Lebiedzinska, M., Szabadkai, G., Jones, A.W., Duszynski, J. and Wieckowski, M.R. (2009). Interactions between the endoplasmic reticulum, mitochondria, plasma membrane and other subcellular organelles. The international journal of biochemistry & cell biology, 41(10), 1805–1816.

Phillips, M.J. and Voeltz, G.K. (2016). Structure and function of ER membrane contact sites with other organelles. Nature reviews Molecular cell biology, 17(2), 69–82.

Denk, W. and Horstmann, H. (2004). Serial block-face scanning electron microscopy to reconstruct three-dimensional tissue nanostructure. PLoS Biol, 2(11), p.e329.

Knott, G., Marchman, H., Wall, D. and Lich, B. (2008). Serial section scanning electron microscopy of adult brain tissue using focused ion beam milling. Journal of Neuroscience, 28(12), 2959–2964.

Briggman, K. L., and Bock, D. D. (2012). Volume electron microscopy for neuronal circuit reconstruction. Current opinion in neurobiology, 22(1), 154–161.

Xiao, C., Chen, X., Li, W., Li, L., Wang, L., Xie, Q. and Han, H. (2018). Automatic mitochondria segmentation for EM data using a 3D supervised convolutional network. Frontiers in neuroanatomy, 12, 92.

Liu, J., Li, L., Yang, Y., Hong, B., Chen, X., Xie, Q. and Han, H. (2020). Automatic reconstruction of mitochondria and endoplasmic reticulum in electron microscopy volumes by deep learning. Frontiers in neuroscience, 14.

Arruda, A. P., Pers, B. M., Parlakgül, G., Güney, E., Inouye, K., and Hotamisligil, G. S. (2014). Chronic enrichment of hepatic endoplasmic reticulum–mitochondria contact leads to mitochondrial dysfunction in obesity. Nature medicine, 20(12), 1427–1435.

McBride, H.M., Neuspiel, M. and Wasiak, S. (2006). Mitochondria: more than just a powerhouse. Current biology, 16(14), R551–R560.

Zhou, Z., Torres, M., Sha, H., Halbrook, C. J., Van den Bergh, F., Reinert, R. B., Yamada, T., Wang, S., Qi, L., et al. (2020). Endoplasmic reticulum–associated degradation regulates mitochondrial dynamics in brown adipocytes. Science, 368(6486), 54–60.

West, M., Zurek, N., Hoenger, A., and Voeltz, G. K. (2011). A 3D analysis of yeast ER structure reveals how ER domains are organized by membrane curvature. Journal of Cell Biology, 193(2), 333–346.

Bruguera, M., Bertran, A., Bombi, J.A. and Rodes, J. (1977). Giant mitochondria in hepatocytes: a diagnostic hint for alcoholic liver disease. Gastroenterology, 73(6), 1383–1387.

Flis, V. V., and Daum, G. (2013). Lipid transport between the endoplasmic reticulum and mitochondria. Cold Spring Harbor perspectives in biology, 5(6), a013235.

Murley, A., Lackner, L.L., Osman, C., West, M., Voeltz, G.K., Walter, P. and Nunnari, J. (2013). ER-associated mitochondrial division links the distribution of mitochondria and mitochondrial DNA in yeast. Elife, 2, e00422.

Lewis, S. C., Uchiyama, L. F., and Nunnari, J. (2016). ER-mitochondria contacts couple mtDNA synthesis with mitochondrial division in human cells. Science, 353(6296).

Wu, H., Carvalho, P., and Voeltz, G. K. (2018). Here, there, and everywhere: The importance of ER membrane contact sites. Science, 361(6401).

Lim, C.Y., Davis, O.B., Shin, H.R., Zhang, J., Berdan, C.A., Jiang, X., Counihan, J.L., Ory, D.S., Nomura, D.K. and Zoncu, R. (2019). ER–lysosome contacts enable cholesterol sensing by mTORC1 and drive aberrant growth signalling in Niemann– Pick type C. Nature cell biology, 21(10), 1206–1218.

Lodhi, I. J., and Semenkovich, C. F. (2014). Peroxisomes: a nexus for lipid metabolism and cellular signaling. Cell metabolism, 19(3), 380–392.

Hernández-Alvarez, M.I., Sebastián, D., Vives, S., Ivanova, S., Bartoccioni, P., Kakimoto, P., Plana, N., Veiga, S.R., Hernández, V., Vasconcelos, N. and Peddinti, G. (2019). Deficient endoplasmic reticulum-mitochondrial phosphatidylserine transfer causes liver disease. Cell, 177(4), 881–895.

Chen, X., Xie, Q., Shen, L. and Han, H. (2018), October. Morphology-Retained Non-Linear Image Registration of Serial Electron Microscopy Sections. In 2018 25th IEEE International Conference on Image Processing (ICIP), 3833–3837. IEEE.

Liu, C., Yuen, J. and Torralba, A. (2010). Sift flow: Dense correspondence across scenes and its applications. IEEE transactions on pattern analysis and machine intelligence, 33(5), 978–994.

Schaefer, S., McPhail, T. and Warren, J. (2006). Image deformation using moving least squares. In ACM SIGGRAPH 2006 Papers, 533–540.

Li, W., Liu, J., Xiao, C., Deng, H., Xie, Q. and Han, H. (2018). A fast forward 3D connection algorithm for mitochondria and synapse segmentations from serial EM images. BioData mining, 11(1), 24.

Stalling, D., Westerhoff, M. and Hege, H.C. (2005). Amira: A highly interactive system for visual data analysis. The visualization handbook, 38, 749–67.

He, K., Zhang, X., Ren, S. and Sun, J. (2016). In Proceedings of the IEEE conference on computer vision and pattern recognition, 770–778.

Chen, L.C., Papandreou, G., Kokkinos, I., Murphy, K. and Yuille, A.L. (2017). Deeplab: Semantic image segmentation with deep convolutional nets, atrous convolution, and fully connected crfs. IEEE transactions on pattern analysis and machine intelligence, 40(4), 834–848.

Chen, L.C., Zhu, Y., Papandreou, G., Schroff, F. and Adam, H. (2018). Encoder-decoder with atrous separable convolution for semantic image segmentation. In Proceedings of the European conference on computer vision (ECCV). 801–818.

Cardona, A., Saalfeld, S., Schindelin, J., Arganda-Carreras, I., Preibisch, S., Longair, M., Tomancak, P., Hartenstein, V. and Douglas, R.J. (2012). TrakEM2 software for neural circuit reconstruction. PloS one, 7(6), e38011.

Keras: Deep learning library for theano and tensorflow, http://keras.io/, 2015.

